# Cross-Attention Over RNA And Protein Sequences Enables Generalizable Interaction Prediction

**DOI:** 10.64898/2026.04.22.720174

**Authors:** Mario Catalano, Gerardo Pepe, Gabriele Ausiello, Romina Appierdo, Claire McWhite, Giorgio Gambosi, Manuela Helmer Citterich, PierFederico Gherardini

## Abstract

Computational predictions are essential to characterize the RNA–protein interaction landscape, yet a persistent gap between benchmark performance and practical utility suggests that current models have limited generalization capabilities. To address this issue, we present CORAL (Cross-attention for RNA-protein Association Learning), a deep learning framework for the prediction of RNA–protein interactions that integrates pretrained protein (ESM-2) and RNA (DNABERT2) language models through bidirectional cross-attention with Low-Rank Adaptation fine-tuning. We also introduce a benchmarking framework that rigorously addresses the problem of data redundancy between training and test sets, which greatly inflates model performances reported in the literature. To this end we adopt three partitioning strategies of increasing stringency: conventional random splits, pairwise non-redundant splits, and component-wise non-redundant splits. CORAL maintains an F1 score of 0.65 under the most stringent component-wise evaluation, compared to 0.47 for the next-best method retaining discriminative behavior. Interpretability analyses further reveal that the cross-attention mechanism captures biologically meaningful features of molecular recognition at two complementary structural scales: at atomic resolution, specific attention heads systematically attend to structurally defined contact positions, showing 26% elevated attention at interface residues across 309 experimentally resolved complexes (*p* < 0.001); at the domain level, protein-side attention localizes to annotated RNA-binding domains across 94 % of 462 proteins examined (median within-protein Cohen’s *d* ≈ 0.94). Together, these findings establish that current RPI prediction benchmarks substantially inflate performance estimates and demonstrate that cross-modal attention architectures yield improved generalization alongside mechanistically interpretable representations.

## Introduction

RNA–protein interactions constitute a fundamental layer of cellular regulation underpinning nearly every aspect of gene expression control ^1–3^. The biological significance of RNA–protein interactions extends beyond normal physiology: dysregulation of RNA-binding proteins has been implicated in diverse pathologies, including neurodegeneration, cancer, and developmental disorders ^4,5^. However, defining the scope of the RNA–protein interactome presents a formidable challenge. The human genome encodes over 1,500 RNA-binding proteins ^6^, while the transcriptome comprises tens of thousands of coding and non-coding RNAs. Even accounting for spatiotemporal constraints such as subcellular compartmentalization and cell-cycle-dependent expression, the combinatorial space of potential interactions remains vast, yet only a fraction has been experimentally characterized. High-throughput methods such as CLIP-seq, RIP-seq, PAR-CLIP, RNAcompete, HITS-CLIP, eCLIP and DIF-FRAC have substantially expanded our knowledge of RNA–protein interactions ^7–13^. However, these approaches require specific antibodies, involve complex experimental protocols, and can exhibit systematic biases that limit coverage^14,15^. Structural methods including X-ray crystallography, nuclear magnetic resonance (NMR) and cryo-electron microscopy provide atomic-resolution insights into binding interfaces but remain low-throughput and inapplicable to many complexes that resist crystallization or present conformational heterogeneity ^16–19^. Computational prediction of RNA–protein interactions offers a complementary approach that can prioritize candidates for experimental validation, enable proteome-wide screening, and generate hypotheses about uncharacterized molecules ^20–22^. The past decade has witnessed remarkable advances in applying deep learning to biological sequence analysis, with transformer-based language models ^23^ emerging as particularly powerful tools for extracting rich representations from evolutionary data ^24^. Protein language models such as ESM2^25^ and ProtTrans^26^, trained on billions of amino acids, have demonstrated the capacity to capture structural and functional information directly from amino acid sequences. Analogous models for nucleic acids, including DNABERT2^27^ and RNA-FM^28^, learn contextual representations that encode sequence patterns relevant to diverse downstream tasks. Several deep learning frameworks have been developed to predict RNA– protein interactions by leveraging these advances in sequence representation. IPMiner employs stacked denoising autoencoders to learn compressed representations from k-mer frequency features, integrating predictions through ensemble learning ^29^. NPI-GNN constructs bipartite interaction networks and applies graph neural networks within the SEAL framework for link prediction, combining network topology with sequence features ^30^. More recent methods have begun to incorporate language model representations: SeqMG processes protein sequences through ESM-2 embeddings fed into graph attention networks, while encoding RNA through multi-scale k-mer features processed by convolutional architectures ^31^. ZHMolGraph similarly utilizes RNA-FM and ProtTrans to generate embeddings that are subsequently integrated with graph neural network features derived from known interaction networks^22^. These methods report strong performance on standard benchmarks, with F1 scores commonly exceeding 0.85. Despite these impressive reported metrics, a troubling disconnect persists between benchmarks and real-world performance, and predictions from different methods often show limited concordance ^32^. This gap suggests that high performance on current evaluation frameworks may not translate to reliable predictions on novel, uncharacterized interactions, precisely the setting where computational methods would provide the greatest value. What accounts for this discrepancy? We propose that the answer lies in how benchmarks are constructed. Standard evaluation protocols partition datasets randomly or with minimal redundancy control, permitting substantial sequence similarity between training and test examples ^33,34^. The problem is particularly acute for interaction prediction, where each data point is a pair of molecules rather than an individual entity ^33^. When training and test sets share similar sequences, models may achieve high benchmark accuracy through memorization rather than by learning generalizable principles of molecular recognition ^35^, and may then fail on the genuinely novel sequences that matter most in practice ^36^. We also identify three architectural limitations in current approaches. First, most methods encode RNA and protein sequences independently and combine them only at the final classification stage, missing the co-dependent features that govern binding; cross-modal attention, which allows each sequence to condition its representation on features of its potential partner and has proven powerful in other multimodal domains ^37–39^, remains largely unexplored for RPI prediction. Second, methods that do incorporate pretrained encoders employ them as frozen feature extractors, although parameter-efficient strategies such as Low-Rank Adaptation (LoRA)^40^ now make end-to-end fine-tuning practical. Third, existing predictors function as black boxes, offering probability scores without insight into the features driving them. In this study, we present CORAL, a deep learning framework that addresses each of these limitations. CORAL integrates pretrained protein and RNA language models, ESM-2 and DNABERT2, respectively, through a bidirectional cross-attention mechanism that explicitly models inter-molecular correspondence during encoding. Unlike previous approaches that employ language models as frozen feature extractors, we fine-tune both encoders using Low-Rank Adaptation, enabling the representations to adapt to the specific requirements of interaction prediction while retaining the knowledge acquired during large-scale pretraining. To rigorously evaluate generalization, we introduce a systematic benchmarking framework comprising three partitioning strategies with increasing stringency: conventional random splits, pairwise non-redundant splits, and component-wise non-redundant splits. We apply this framework to benchmark CORAL against four state-of-the-art methods (IPMiner, NPI-GNN, SeqMG, and ZHMolGraph) revealing that performance under stringent conditions diverges dramatically from conventional benchmarks. Finally, we conduct interpretability analyses demonstrating that specific attention heads in our cross-modal architecture learn to attend to structurally defined contact positions between RNA and protein molecules, providing evidence that the model captures biologically meaningful features of molecular recognition. Together, these contributions advance both the practical reliability and mechanistic transparency of computational RNA–protein interaction prediction.

## Results

### Overview of the CORAL Architecture

To address the limitations of existing RPI predictors, we developed CORAL (Cross-attention for RNA-protein Association Learning), a dual-encoder deep learning framework that couples pretrained biological language models with an explicit cross-modal fusion mechanism (Figure 1). RNA and protein sequences are first encoded independently by DNABERT2 and ESM2-150M, respectively, yielding contextualized token-level embeddings that capture modality-specific biochemical and evolutionary features. Because the two encoders operate in spaces of different dimensionality, protein token embeddings are projected from 640 to 768 dimensions to match the RNA representation space prior to fusion. The aligned embeddings are then passed through a cross-modality encoder composed of two stacked layers, each integrating bidirectional cross-attention, enabling simultaneous RNA-to-protein and protein-to-RNA information exchange, with modality-specific self-attention and position-wise feed-forward networks. This design allows each sequence to selectively attend to relevant features in its partner while preserving its own contextual structure, in contrast to late-fusion approaches that concatenate independently learned representations. The refined token embeddings are aggregated by masked mean pooling, concatenated into a joint representation, and fed into a linear classification head that outputs the interaction probability. Both language model backbones are fine-tuned jointly with the cross-modality encoder through Low-Rank Adaptation (LoRA), enabling end-to-end optimization while keeping the number of trainable parameters tractable. A full specification of the architecture and training procedure is provided in Materials and Methods.

**Figure 1.**
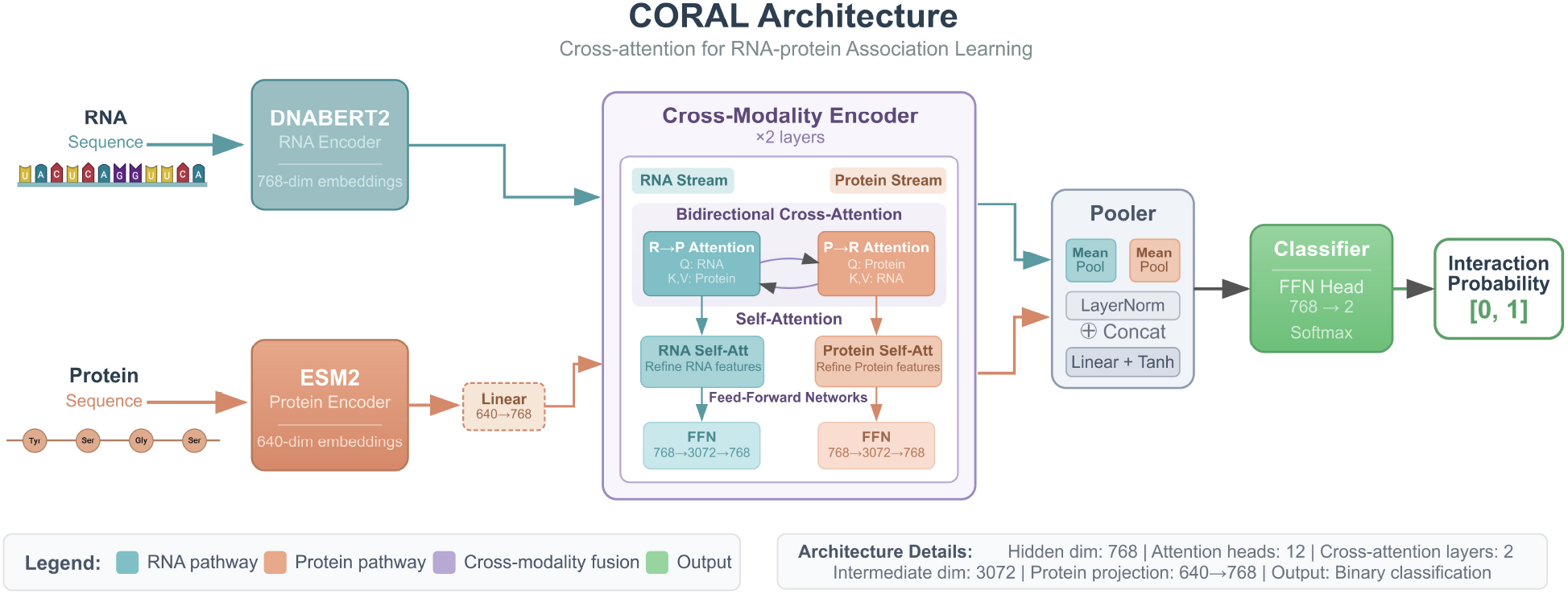
Overview of CORAL Architecture. The model employs a dual-encoder framework with pretrained language models: DNABERT2 for RNA sequences and ESM2 for protein sequences. Protein embeddings are projected from 640 to 768 dimensions to match the RNA embedding space. Both representations are then processed through a cross-modality encoder consisting of two stacked layers, each comprising bidirectional cross-attention (enabling RNA-to-protein and protein-to-RNA information exchange), modality-specific self-attention, and feed-forward networks. The refined embeddings are aggregated via mean pooling with layer normalization, concatenated, and passed through a classification head to predict interaction probability. This architecture enables the model to learn rich cross-modal representations by allowing each modality to attend to relevant features in the other while preserving modality-specific contextual information.

### Dataset Compilation and Quality Control

To obtain sufficient data for model training and validation, we compiled RNA–protein interactions from five publicly available datasets: RNAInter, RPI488, RPI369, RPI2241, and RPI1807^29,41–43^. The datasets with the RPI prefix comprise RNA-Protein interactions derived from experimentally determined PDB complexes, while RNAInter provides interactions curated from diverse sources, including high-throughput experiments, low-throughput experiments, and computational predictions. To ensure data quality, we applied stringent selection criteria. From RNAInter, only interactions categorized as having “strong experimental evidence” were retained. From the RPI datasets, exclusively positive interactions were selected. Additionally, we excluded proteins exceeding 1,250 amino acid residues and RNAs exceeding 6,000 nucleotides in length, constraints imposed by GPU memory limitations during model training. These filtering criteria yielded 24,012 high-confidence positive interactions.

### Two redundancy criteria define a stringency ladder for RPI benchmarking

Standard random train–test splits can substantially overestimate the generalization capacity of RPI prediction models, as test pairs often retain high sequence similarity to training examples. To rigorously evaluate the generalization capacity of RPI prediction models, we propose two distinct operational definitions of redundancy between training and test sets, differing in their stringency. The first definition, termed pairwise redundancy, classifies two RPI pairs as redundant only if both the RNA and protein components simultaneously exceed specified sequence identity thresholds. Under this criterion, an RPI pair in the test set may share sequence similarity with training pairs in one component (either RNA or protein) but not in both components simultaneously (Figure 2a). The second definition, termed component-wise redundancy, classifies two RPI pairs as redundant if either the RNA or protein component exceeds the specified sequence identity thresholds. This more stringent criterion ensures that no individual RNA or protein sequence in the test set is closely related to any sequence in the training set, irrespective of its interaction partner (Figure 2a). A worked example illustrating how each definition assigns pairs to training and test sets is shown in Figure 2b. Together, these complementary criteria form the basis for the train–test partitioning strategies described below.

**Figure 2.**
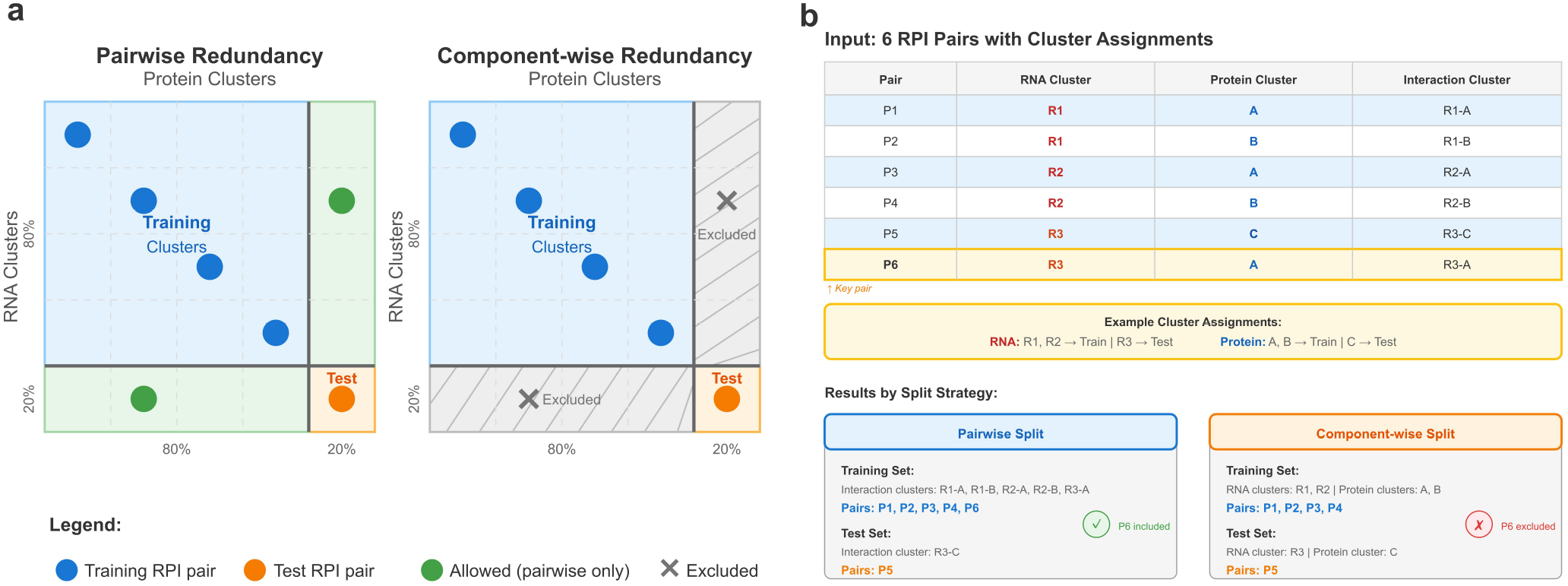
Redundancy definitions for training-test partitioning of RNA-protein interaction datasets. **(a)** Two operational definitions of redundancy between RPI pairs are illustrated: pairwise redundancy, which requires simultaneous sequence similarity in both RNA and protein components, and component-wise redundancy, which requires sequence similarity in either component. Under pairwise redundancy (left), test set pairs may share cluster membership with training pairs in one component (green regions), provided they differ in the other. Under component-wise redundancy (right), pairs with mixed cluster assignments—where one component belongs to a training cluster and the other to a test cluster—are excluded (gray hatched regions), ensuring complete separation at the individual component level. **(b)** Worked example demonstrating the practical consequences of each redundancy definition. Six RPI pairs (P1–P6) with their corresponding RNA and protein cluster assignments are shown. Given the example cluster assignments (RNA: R1, R2 → training, R3 → test; Protein: A, B → training, C → test), pair P6 (RNA cluster R3, Protein cluster A) is included in the training set under pairwise splitting, as its interaction cluster (R3-A) differs from the test interaction cluster (R3-C). Conversely, under component-wise splitting, P6 is excluded due to its mixed cluster assignment.

### Construction of Train-Test splits with progressively increasing stringency

Building on the aforementioned redundancy definitions, we generated three families of train-test partitions of increasing stringency (see Materials and Methods for full procedures). The first, a conventional random split (hereafter *redundant* split), imposes no constraint on sequence similarity and therefore produces overlapping training and test subsets. The second, the *pairwise* split, excludes from the test set any pair whose RNA and protein components both exceed their respective identity thresholds relative to training pairs; high similarity in a single component is still permitted. The third and most stringent, the *component-wise* split, ensures that no individual RNA or protein sequence exceeding its identity threshold is shared between training and test sets, regardless of interaction partner. Both the pairwise and component-wise splits were constructed at two levels of stringency, using a permissive threshold pair (80% identity for RNA, 60% for protein) and a stricter pair (50% for RNA, 30% for protein). To mitigate sampling bias, five independent replicates were generated for each strategy and threshold combination, yielding ten pairwise and ten component-wise datasets in addition to five redundant datasets. As expected, the component-wise criterion proved the most restrictive and retained the fewest pairs (Table 1). For each partition, balanced datasets were then assembled by generating negative (non-interacting) pairs: RNA and protein sequences were randomly re-paired within the corresponding training or test subset, and any pair coinciding with a known interaction was discarded, so that negative samples represented putative non-interactions. All resulting datasets contained equal numbers of positive and negative samples (Table 1), enabling systematic evaluation of model performance under progressively more demanding generalization conditions.

**Table 1.** Summary of dataset partitions for model training and evaluation. Values represent the range (minimum–maximum) across five independent folds. The component-wise and pairwise splits enforce sequence identity thresholds to minimize data leakage, while the redundant split serves as a baseline without such constraints. All datasets maintain equal distribution between positive (interacting) and negative (non-interacting) RNA-protein pairs.

| Split Strategy | Seq. Identity Threshold (RNA / Protein) | Training Samples | Test Samples |
| --- | --- | --- | --- |
| Component-wise | 80% / 60% | 22,110–24,100 | 3,744–4,264 |
| Pairwise | 80% / 60% | 38,154–38,202 | 9,822–9,870 |
| Component-wise | 50% / 30% | 22,056–22,624 | 4,156–4,330 |
| Pairwise | 50% / 30% | 38,134–38,190 | 9,834–9,890 |
| Redundant | — | 42,468 | 10,618 |

### CORAL Exhibits Superior Generalization to Novel RNA-Protein Pairs

We benchmarked our model (CORAL) against four state-of-the-art RPI prediction models: IPMiner, NPI-GNN, SeqMG, and ZHMolGraph. All models were trained and evaluated from scratch on each of the three partition strategies. Figure 3 presents the average performance metrics on the redundant split, pair-wise split, and component-wise split test sets using sequence identity thresholds of 80% for RNA and 60% for protein. On the redundant split, all models achieved strong performance, with CORAL attaining the highest F1 score (0.92) and accuracy scores (0.92). Performance on the pairwise split remained relatively high, with only moderate degradation observed across all models: CORAL maintained an F1 score of 0.91, while other models exhibited F1 scores ranging from 0.75 to 0.89. In stark contrast, all competing models exhibited substantial performance degradation on the component-wise split, revealing limited generalization to sequences not seen during training. CORAL was the only model to maintain discriminative predictions while achieving a meaningful F1 score, reaching 0.65 on the 80/60 split, with ZHMolGraph, SeqMG and IPMiner attaining F1 scores of 0.47, 0.43 and 0.03, respectively. NPI-GNN produced an F1 of 0.67 on the same split, which superficially appears comparable to CORAL, but inspection of its component metrics reveals that this score is a degenerate artifact rather than evidence of generalization. NPI-GNN classified 99% of test pairs as interacting regardless of sequence content (recall = 0.99, specificity = 0.03); on a dataset with 50:50 class balance, such indiscriminate positive predictions mechanically yield precision = 0.50 and recall = 1.00, forcing F1 toward 0.67 by construction. IPMiner exhibited the mirror failure mode, predicting nearly every pair as non-interacting (recall = 0.01, specificity = 0.99) and collapsing F1 to 0.03. Both models therefore failed to learn any discriminative signal under stringent conditions and defaulted to single-class prediction, with their apparent F1 values determined entirely by the direction of collapse and the class balance of the test set rather than by any residual predictive ability. This pattern intensified under stricter identity thresholds (50% RNA, 30% protein), where all methods degraded further on the component-wise split (Supplementary Figure 1). CORAL consistently attained the highest F1 and accuracy across splitting strategies and thresholds, demonstrating superior generalization capabilities.

**Figure 3.**
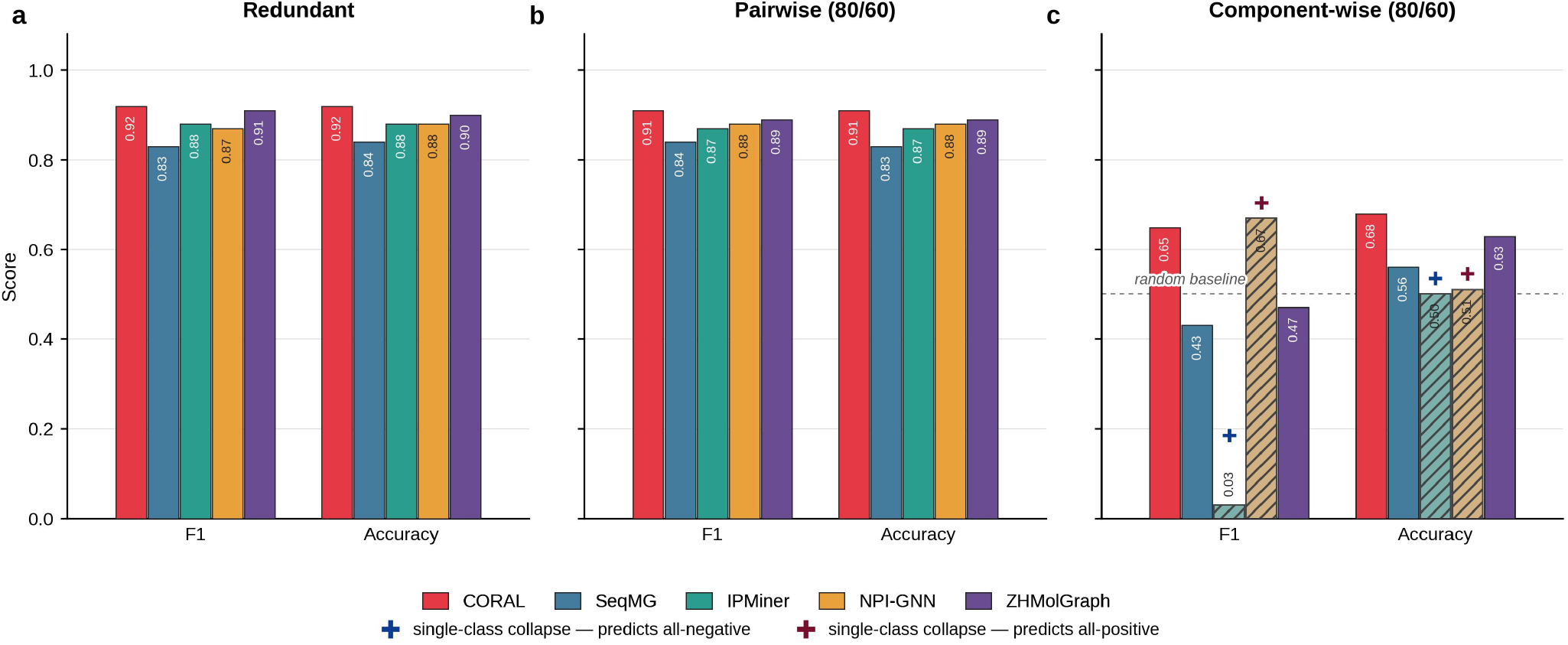
Performance comparison of CORAL against baseline methods across data partitioning strategies of increasing stringency. F1 and Accuracy are shown for five RNA–protein interaction predictors: CORAL (this work, red), SeqMG (blue), IPMiner (teal), NPI-GNN (amber) and ZHMolGraph (purple) on **(a)** the conventional redundant split, **(b)** the pairwise (80/60) non-redundant split, and **(c)** the component-wise (80/60) non-redundant split. Numbers in parentheses denote the sequence identity thresholds (RNA / protein) used to construct the pairwise and component-wise partitions; the redundant split is threshold-independent. Bar heights are means over five independent train/test folds, and the exact value is annotated on each bar. CORAL attains the highest F1 and Accuracy under all three partitioning regimes. The dashed grey reference line in (c) marks the F1 and Accuracy expected from a random predictor on the class-balanced (50:50 positives:negatives) test set. Bars marked with a cross identify methods that collapse to single-class prediction on the component-wise split; for these methods, F1, Recall and Specificity are mechanically determined by the direction of collapse and the class balance and therefore do not reflect discriminative ability. Cross colour encodes the direction of collapse: navy denotes “predicts all-negative”, whereas maroon denotes “predicts all-positive”. Diagonal hatching and desaturated colour of the affected bars visually reinforce these crosses.

### Encoder Fine-Tuning is Critical for Generalization to Novel Sequences

To the best of our knowledge, existing RPI prediction methods that incorporate pretrained protein and RNA language models, such as SeqMG and ZHMolGraph, employ these encoders as frozen feature extractors, training only the downstream components on interaction data. To assess the contribution of end-to-end encoder adaptation in CORAL, we conducted an ablation study comparing two training regimes (Figure 4): one in which both the ESM2 and DNABERT2 weights were kept frozen during training, and one in which they were jointly fine-tuned with LoRA alongside the cross-modality encoder and the classification head. Both variants were trained and evaluated on the redundant, pairwise (80% RNA / 60% protein), and component-wise (80% RNA / 60% protein) splits under matched optimisation settings, with the initial learning rate adjusted for each regime to account for the difference in the number and initialisation state of the trainable parameters (see Methods). On the redundant split, fine-tuning produced a modest but consistent gain, increasing both the F1 score and the accuracy from 0.89 to 0.92 (+0.03, Figure 4a). A comparable improvement was observed on the pairwise split, where the F1 score and the accuracy rose from 0.87 to 0.91 (+0.04, Figure 4b). The most pronounced effect emerged on the component-wise split, the most stringent evaluation setting: LoRA fine-tuning increased the F1 score from 0.53 to 0.65 (+0.12) and the accuracy from 0.63 to 0.68 (+0.05, Figure 4c). The gain from LoRA fine-tuning was therefore largest on the most stringent split, where the relative improvement in F1 score was approximately four-fold larger than on the redundant split.

**Figure 4.**
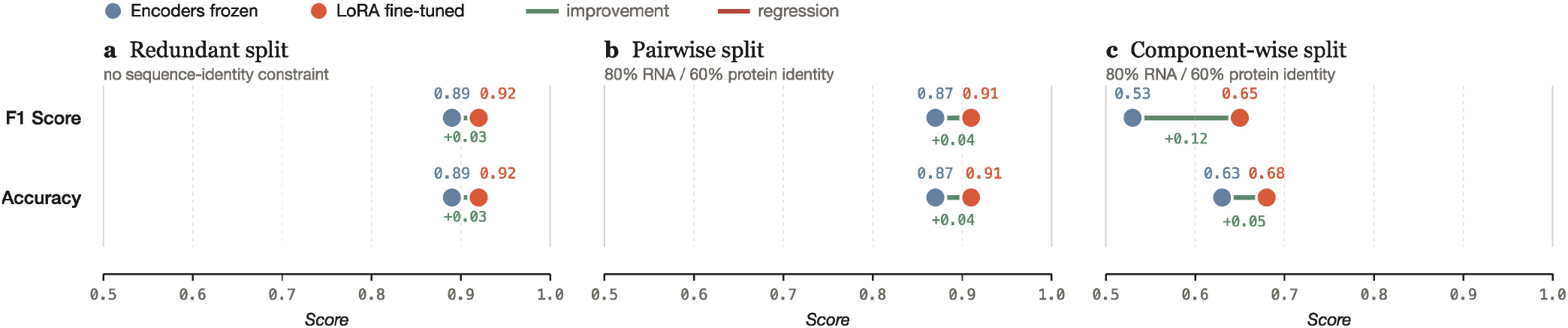
End-to-end encoder fine-tuning with LoRA is critical for model generalization to novel sequences. Dumbbell plots comparing the predictive performance of CORAL when employing pretrained language models as frozen feature extractors (blue dots) versus jointly fine-tuning the encoders using Low-Rank Adaptation (red dots). Performance metrics (F1 score and accuracy) are evaluated across three data-partitioning strategies of increasing stringency: **(a)** Redundant split with no sequence identity constraints, **(b)** Pairwise split enforcing 80% RNA and 60% protein sequence identity thresholds, and **(c)** Component-wise split utilizing the same 80%/60% thresholds. Green solid lines indicate the absolute performance improvement achieved by the fine-tuned variant. While LoRA fine-tuning provides modest but consistent gains under redundant and pairwise evaluation conditions, it drives a substantial performance increase on the most stringent component-wise split (F1 score +0.12).

### Cross-Attention Mechanism Learns Structural Contact Interfaces

To investigate the attention mechanisms underlying CORAL’s interaction predictions, we conducted an explainability analysis focused on correctly classified interacting pairs. We focused on the 415 RNA-protein complexes in the PDB with resolution ≤ 3.5 Å that (1) represent true interactions with known interface residues derived from the co-crystallized complex (see Methods for proximity threshold), and (2) were correctly predicted as “interacting” by CORAL. This focus on true positive predictions allows us to examine what features the model attends to when making accurate interaction calls. For each complex, we defined ground-truth structural contacts at the residue level: a nucleotide-amino acid pair was classified as a “contact” if any inter-atomic distance was ≤ 3.4 Å in the experimental structure. To ensure sufficient contact density and sequence length for robust statistical analysis, we applied quality filters requiring (1) a minimum of three contacts per complex, and (2) minimum sequence lengths of 15 nucleotides (RNA) and 25 amino acids (protein). These filters yielded a final dataset of 309 complexes with well-defined structural contact annotations. We extracted cross-attention scores from CORAL for all 309 complexes and examined whether attention scores were systematically elevated at structurally defined contact positions compared to non-contact positions. To identify which attention heads exhibited differential contact-position responses while controlling for multiple comparisons, we first conducted a Friedman test across all 12 heads, comparing per-complex mean differences (Δ_*i*_ = mean contact attention − mean non-contact attention) across the 309 complexes. This test revealed highly significant variation across heads (*χ*^2^(11) = 207, *p* << 0.001; Figure 5c). We then performed post-hoc pairwise Wilcoxon signed-rank tests comparing the head with the largest median Δ against each of the remaining heads, with Benjamini–Hochberg false discovery rate (FDR) correction (*α* = 0.05) to identify which specific heads exhibited elevated contact attention. Head 7 emerged as the head with the largest median contact bias. Head 0 was retained alongside head 7 as it was not statistically distinguishable from it (Wilcoxon adj-*p* = 0.55), whereas the remaining ten heads all exhibited significantly lower contact bias (adj-*p* ≤ 4 × 10^−6^; Figure 5c). All subsequent analyses focused on attention scores averaged across these two screened heads. The spatial coincidence between elevated attention and annotated contacts is illustrated by three exemplar complexes spanning distinct recognition logics: Roquin-1 ROQ in complex with the Hmgxb3 CDE stem-loop (PDB 4QIL, winged-helix fold, 14 contacts, per-complex Cohen’s *d* = 1.51), the RRM-fold C-terminal RNA-binding domain of the Bacillus subtilis DEAD-box helicase YxiN in complex with a fragment of 23S ribosomal RNA spanning hairpin 92 of the peptidyl-transferase center (PDB 3MOJ, RRM fold, 17 contacts, *d* = 0.99), and the SelB C-terminal domain in complex with its SECIS hairpin (PDB 2UWM, tandem winged-helix array, 9 contacts, *d* = 1.56) (Figure 5a). Projecting the average of heads 0 and 7 attention onto the three-dimensional structure of the 4QIL complex, with grey lines connecting protein C*α* atoms and central RNA nucleotides for cells in the top 1% of the attention matrix, shows that high-attention residues localize to the winged-helix face contacting the CDE stem-loop and that the top-1% lines converge on the apical tri-loop and closing base pair of the CDE stem (Figure 5b). To rigorously assess whether heads 0 and 7 systematically prioritize structural contact positions, we employed three complementary statistical approaches that account for the hierarchical structure of our data (positions nested within complexes) and the severe class imbalance (contact positions representing *<*1% of all possible interactions, with an average ratio of 1:270), avoiding pseudoreplication that could artificially inflate significance. First, treating each complex as an independent unit of analysis to avoid pseudoreplication, we calculated the mean difference between contact and non-contact attention scores for each of the 309 complexes (Δ_*i*_ = mean contact attention − mean non-contact attention). The population-level analysis revealed that attention scores at contact positions (mean = 0.0250, SD = 0.0142) were significantly elevated relative to non-contact positions (mean = 0.0198, SD = 0.0091), representing a 26% relative increase (mean Δ = 0.00512; one-sample *t*-test: *t*(308) = 9.8, *p <* 0.001). This difference was remarkably consistent, with 244 of 309 complexes (79%) exhibiting positive differences (sign test: *p* < 0.001). Second, to validate these findings under relaxed distributional assumptions, we performed hierarchical bootstrap resampling with 10,000 iterations, resampling complexes (rather than individual positions) with replacement. This procedure yielded a bootstrap standard error of 0.00052 and 95% confidence interval of [0.0041, 0.0062] that excluded zero. Third, to assess the magnitude of this effect within individual complexes, we computed Cohen’s *d* independently for each complex. The resulting distribution of effect sizes had a mean of 0.35, a median of 0.31, and a standard deviation of 0.49, with 79% of complexes showing positive effect sizes (Figure 5d, blue curve). Collectively, these three analytical approaches provide robust statistical evidence that the attention scores of heads 0 and 7 are systematically higher for structural contact positions. To assess whether this contact-position bias represents a property specific to heads 0 and 7 or a feature distributed across the attention mechanism, we repeated all three analyses using attention scores averaged across all 12 heads. Under this regime, the mean per-complex difference decreased from Δ = 0.00512 to Δ = 0.00175 (66% attenuation), the mean Cohen’s *d* decreased from 0.35 to 0.17 (~ 53% attenuation), the median Cohen’s *d* decreased from 0.31 to 0.13 (~ 58% attenuation), and the proportion of complexes with positive bias decreased from 79% to 61% (Figure 5d, grey curve). Importantly, the all-heads average remained significantly contact-biased: 190 of 309 complexes (61%) showed a positive difference (sign test *p <* 0.001), the one-sample t-test remained highly significant (*t* = 6.5, *p* < 0.001), and the hierarchical-bootstrap 95% confidence interval [0.0012, 0.0023] excluded zero. By comparison, under the screened average of heads 0 and 7 only 65 of 309 complexes (21%) showed Cohen’s *d* ≤ 0. The contact-position signal is therefore distributed across the attention mechanism but concentrated in heads 0 and 7, rather than being a near-exclusive property of those two heads.

**Figure 5.**
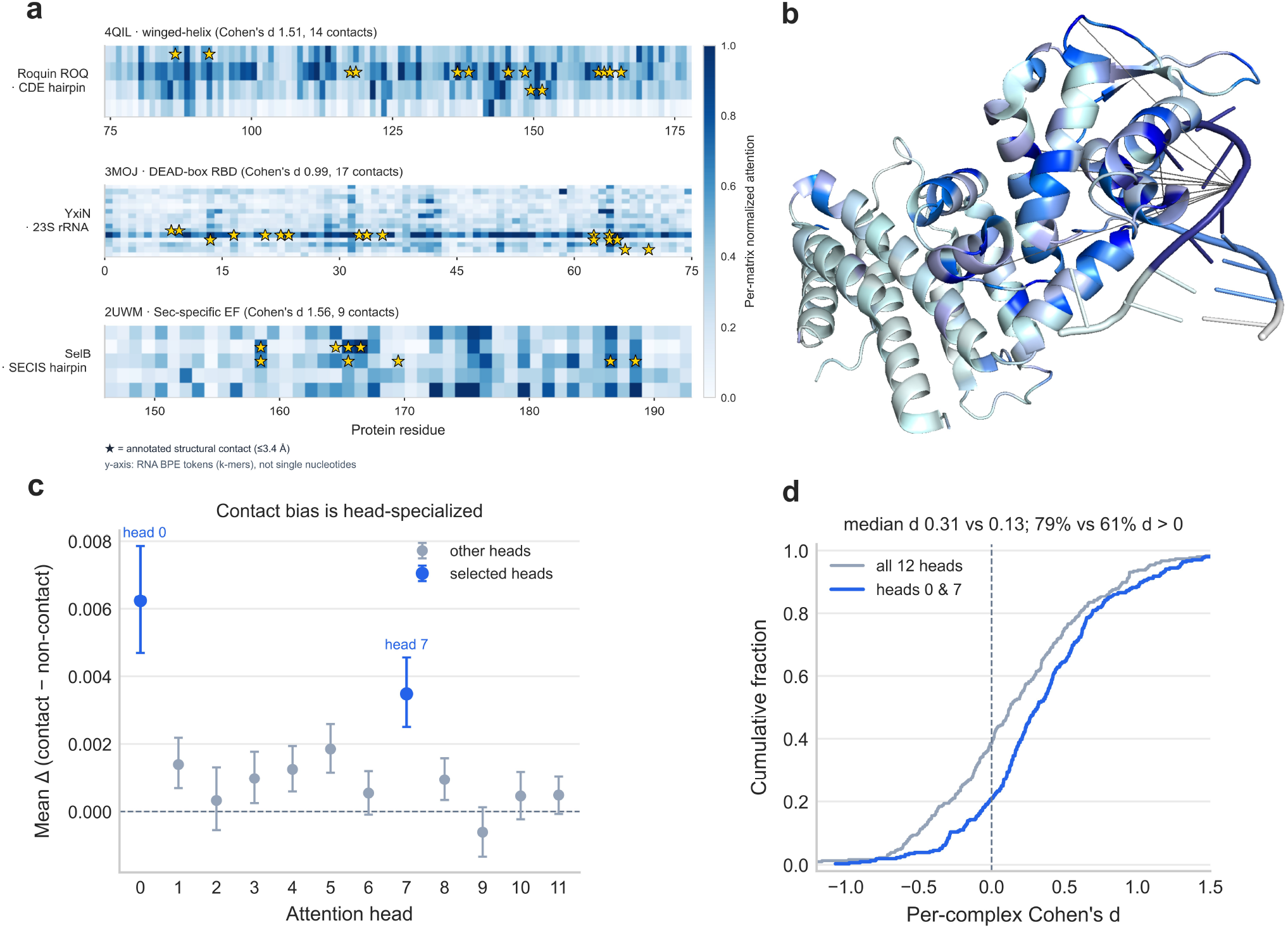
CORAL’s cross-attention is elevated at structurally defined contact positions across 309 RNA–protein complexes. **(a)** Per-complex cross-attention heatmaps averaged across the two screened heads (0 and 7), for three exemplar complexes spanning distinct recognition logics: Roquin-1 ROQ in complex with the Hmgxb3 CDE stem-loop (PDB 4QIL, winged-helix fold), the RRM-fold C-terminal RNA-binding domain of the Bacillus subtilis DEAD-box helicase YxiN in complex with a fragment of 23S ribosomal RNA spanning hairpin 92 of the peptidyl-transferase center (PDB 3MOJ, RRM fold), and the SelB C-terminal domain in complex with its SECIS hairpin (PDB 2UWM, tandem winged-helix array). Rows are RNA byte-pair-encoding tokens (DNABERT2 k-mers, not single nucleotides); columns are protein residues, cropped to the contact-spanning window *±* 12 residues. Each matrix is independently min–max normalized at its 2nd–98^th^ percentile (white → deep-blue colormap, shared colorbar). Gold stars denote annotated structural contacts, defined as nucleotide–amino acid pairs with any inter-atomic distance ≤ 3.4 Å in the experimentally determined PDB structure. Subplot titles report the per-complex Cohen’s *d* of the contact-versus-non-contact attention distribution and the number of contacts. **(b)** Three-dimensional structure of the 4QIL Roquin-ROQ CDE hairpin complex, with protein residues and RNA nucleotides coloured by their maximum cross-attention score along the partner axis (white, low; deep blue, high) under per-molecule scaling. Grey lines connect protein C*α* atoms to the central nucleotide of any RNA k-mer token whose attention falls within the top 1% of the full attention matrix. **(c)** Per-head contact bias across the 12 cross-attention heads, computed as Δ = mean(attention at contact cells) − mean(attention at non-contact cells) per complex and averaged across complexes. Points denote means; whiskers denote 95% confidence intervals from a hierarchical bootstrap with 5,000 iterations resampling whole complexes (rather than individual cells), so the interval reflects between-complex variability. Heads 0 and 7 are shown in deep blue; the remaining ten heads, which exhibited significantly lower bias than head 7 (adj-*p* ≤ 4 *×* 10^−6^), are shown in grey. Head 0 shows a larger mean Δ than head 7 in this panel, but head 7 has the larger median Δ and was taken as the reference head for the post-hoc comparison; head 0 was retained because it was not statistically distinguishable from head 7 (adj-*p* = 0.55). The dashed grey line marks Δ = 0. **(d)** Empirical cumulative distribution of the per-complex Cohen’s *d* under two head-aggregation schemes: the screened subset of heads 0 and 7 (blue, heads_best) and the simple average across all 12 heads (grey, heads_all). The dashed vertical line marks *d* = 0. Annotations above the panel report the median *d* under each scheme (0.31 versus 0.13) and the proportion of complexes with *d >* 0 (79% versus 61%).

### Protein-Side Attention Localizes to Annotated RNA-Binding Domains

Having established that specific cross-attention heads attend to atomic contact positions in experimentally resolved RNA–protein complexes, we next examined whether the cross-attention mechanism also recapitulates structure at a coarser, sequence-domain resolution on the protein side. To this end, we conducted a parallel analysis testing whether CORAL’s RNA-to-protein cross-attention, marginalized over the RNA axis, is systematically elevated at residues annotated as canonical RNA-binding domains (RBDs). We first assembled a protein universe comprising all proteins that appear as the positive partner in any interaction within our curated dataset and annotated them against Pfam using an RBD whitelist drawn from the human RBP census of Gerstberger et al. ^6^. The whitelist was constructed by retaining the domain families observed in at least four distinct RBPs in that census, which yielded 85 RBD families encompassing canonical sequence-specific RBDs (RRM, KH, dsRBD, CCCH and CCHC zinc fingers, CSD, PUF, Piwi, YTH, La, Sm/LSM, S1, among others) as well as RNA-associated modules including ribosomal proteins, translation factors, helicases, and RNA-modification enzymes. Of 3,814 proteins examined, 624 carried at least one annotated RBD. We further required that each protein exceed 100 amino acid residues in length so that sufficient non-RBD sequence existed for comparison, exhibit RBD coverage not exceeding 70% of its length so that the inside-versus-outside contrast remained informative, and have at least one true-positive RNA partner (a curated-positive interaction predicted by CORAL with probability greater than 0.5). These criteria yielded 462 proteins aggregated from 12,834 true-positive (protein, RNA) pairs, with the per-protein partner count following a strongly right-skewed distribution (median = 1, mean = 27.8, maximum = 825). For each (protein, RNA) pair, we extracted the RNA-to-protein cross-attention map from the final cross-modality layer, averaged it over the selected attention head(s), and marginalized over the RNA axis by mean reduction to obtain a per-residue protein attention score. Per-pair vectors were then averaged element-wise across each protein’s true-positive partners to produce a per-protein consensus attention profile, ensuring that the protein constituted the independent unit of analysis. For each protein, we computed Δ_*k*_ = mean(*s* | residue ∈ any RBD) − mean(*s* | residue ∉ any RBD), the difference between mean attention at residues falling within annotated RBD intervals and at residues outside them, where the inside set is defined as the union of all of a protein’s annotated RBD intervals. To identify which of the 12 attention heads exhibit differential responses at RBD residues while controlling for multiple comparisons, we first conducted a Friedman test on the per-protein Δ_*k*_ values across the 12 heads, which revealed highly significant variation (*χ*^2^(11) = 429.08, *p <<* 0.001). Following this result, we performed post-hoc pairwise Wilcoxon signed-rank tests comparing the head with the largest median Δ_*k*_ against each of the remaining heads, with Benjamini–Hochberg false discovery rate correction (*α* = 0.05). Head 2 emerged as having significantly higher RBD-biased attention than every other head (all eleven adjusted *p* − *values <<* 0.001) and was retained as the sole selected head for subsequent analyses, notably distinct from heads 0 and 7, which were selected by the atomic-contact screen. To rigorously assess whether head 2 systematically prioritizes RBD residues, we applied four complementary statistical approaches mirroring the contact-analysis pipeline, treating each protein as an independent unit to avoid pseudoreplication. First, a one-sample *t*-test on the population of {Δ_*k*_} values confirmed that the mean per-protein difference significantly exceeded zero (*t*(461) = 27.08, *p* ≪ 0.001; mean(Δ) = 2.95 *×* 10^−3^ attention units, SD = 2.34 *×* 10^−3^; Figure 6b). Second, the non-parametric sign test revealed that 435 of 462 proteins (94.2%) exhibited Δ_*k*_ *>* 0 (exact one-sided binomial *p* ≪ 0.001), demonstrating that the population-level effect reflects a near-universal directional pattern rather than one driven by a few large-magnitude outliers. Third, hierarchical bootstrap resampling with 10,000 iterations, resampling proteins (rather than individual residues) with replacement, yielded a bootstrap standard error of 1.09 *×* 10^−4^ and a 95% confidence interval of [2.74 *×* 10^−3^, 3.17 *×* 10^−3^] that excluded zero by a wide margin. Fourth, the distribution of per-protein Cohen’s *d* effect sizes, a within-protein standardized measure of the inside-versus-outside separation, was strongly right-shifted (mean(*d*) = 0.88, median(*d*) = 0.94, SD = 0.48), with 435 of 462 proteins (94.2%) exhibiting positive effects (Figure 6c). Representative consensus attention profiles for five proteins spanning distinct RBD architectures — CELF1 (three RRM domains), YTHDF1 (one YTH domain), ZFP36 (two zf-CCCH zinc fingers), STAU1 (three dsRBD domains), and KHSRP (four KH domains), illustrate a strong spatial coincidence between regions of elevated attention and annotated RBDs (Figure 6a). To rule out trivial positional confounds, such as attention being systematically higher in the middle of the protein sequence, we performed a position-shuffle null analysis in which, for each protein, the RBD residue mask was cyclically rotated along the sequence by a random non-zero offset, preserving the number, lengths, and gap structure of the intervals while randomizing their absolute placement. Recomputing the aggregate Δ over 1,000 random rotations yielded a null distribution centered near zero (mean = −1.0 *×* 10^−5^, SD = 8.7 *×* 10^−5^, maximum = 2.86 *×* 10^−4^), well below the observed value of 2.95 *×* 10^−3^. The observed effect exceeded all 1,000 null draws (empirical *p <* 0.001) and lay approximately 34 standard deviations above the null mean (Figure 6b). To determine whether the population-level signal was driven by a single dominant RBD class or generalized across diverse RBD families, we stratified the proteins by their dominant annotated domain and computed family-specific effect sizes for the 14 families containing at least 10 proteins, with 95% confidence intervals obtained from 2,000 bootstrap resamples. The across-protein effect was positive and statistically significant in 13 of the 14 stratified families, spanning canonical sequence-specific RBDs (RRM with *n* = 107, *d* = 1.68; KH with *n* = 37, *d* = 2.15; CSD with *n* = 18, *d* = 2.51; both C2H2 and CCCH classes of zinc finger) and RNA-associated modules (KOW, S4, LSM, DEAD, Piwi, MMR_HSR1, and GTP_EFTU), with effect sizes ranging from *d* = 0.89 (GTP_EFTU) to *d* = 6.32 (KOW). A single family ran in the opposite direction: the PUA domain (*n* = 11) exhibited a negative effect size (*d* = −0.72; 95% CI [−1.42, −0.26]) (Figure 6d). PUA domains commonly occur as accessory modules in RNA-modification enzymes (e.g., pseudouridine synthases, tRNA-modification proteins) where the primary RNA-contacting residues lie outside the annotated PUA interval, on catalytic regions captured by the “outside” set. Given *n* = 11, this should be regarded as an anomalous family rather than a robust counter-pattern.

**Figure 6.**
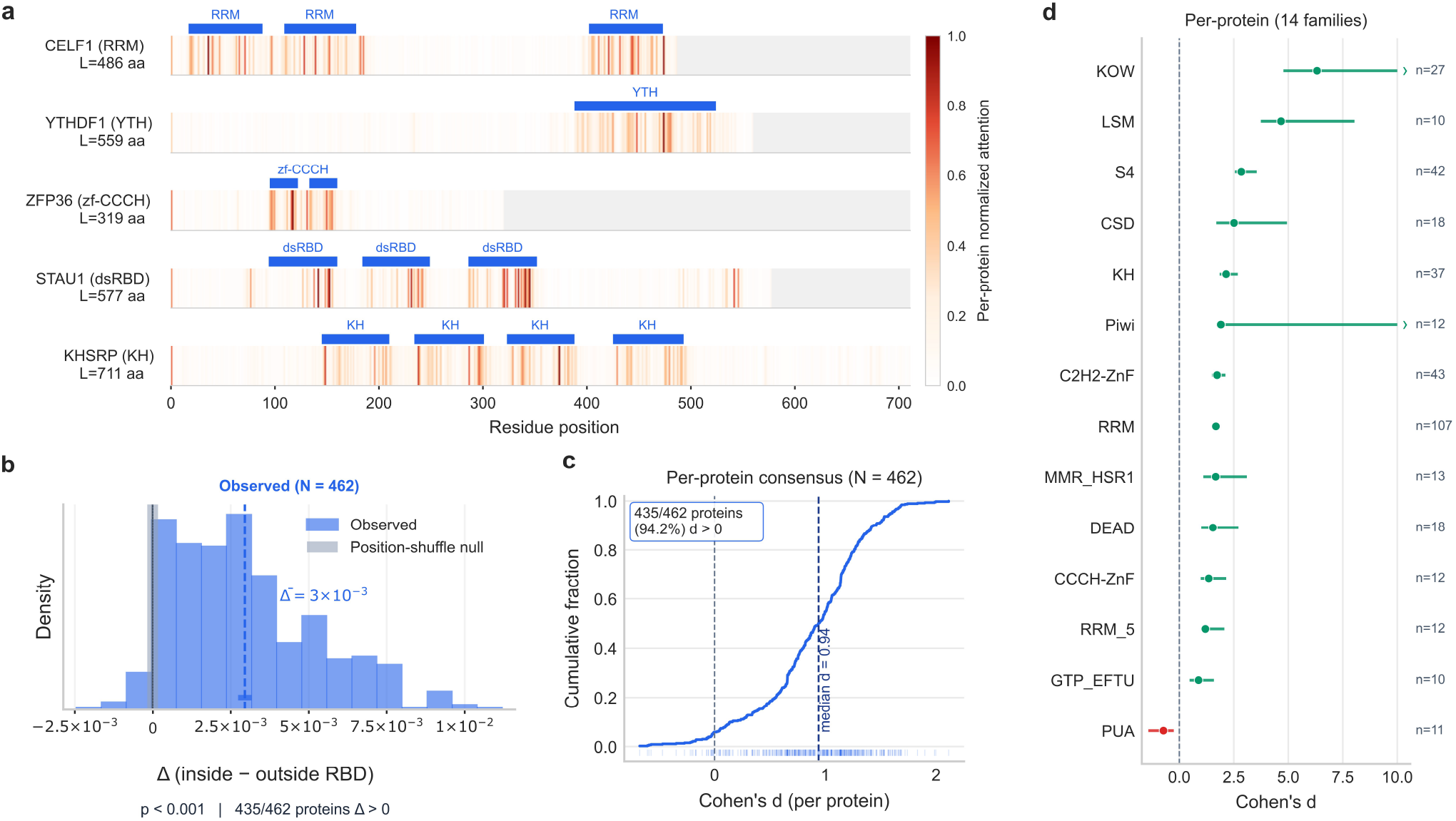
Per-protein cross-attention scores from CORAL at annotated RNA-binding domains across 462 proteins. **(a)** Per-residue consensus cross-attention scores along the protein sequence for five representative RNA-binding proteins spanning distinct RBD architectures. For each protein, the consensus score at a given residue is the protein-marginalized cross-attention from the selected head of the final cross-modality layer, averaged across the protein’s true-positive RNA partners, and normalized to the [0, 1] range within each protein for visualization (white, low; dark red, high). Grey regions correspond to residue positions beyond the protein’s length. Blue bars above each track indicate the boundaries of the annotated RBD intervals retrieved from Pfam, labeled by domain family. **(b)** Distribution of the per-protein inside-versus-outside RBD attention difference across the 462 cohort proteins (blue), shown alongside the position-shuffle null distribution generated from 1,000 random cyclic rotations of the per-protein RBD masks along the sequence (grey band); the rotation preserves the number, lengths, and gap structure of the annotated intervals while randomizing their placement. Because the null is far more tightly concentrated than the observed distribution, it is shown not as a separate histogram but as a shaded grey band spanning its central 95% interval at zero. The dashed blue line marks the observed mean. Annotations below the panel report the empirical p-value of the position-shuffle null and the number of proteins with Δ_*k*_ *>* 0. **(c)** Empirical cumulative distribution function of the per-protein within-protein Cohen’s d across the 462 proteins, with population variances. The dashed vertical line indicates the median of the distribution; the annotation reports the fraction of proteins with positive effect size. Tick marks along the bottom show the individual per-protein values. **(d)** Forest plot of the across-protein effect size computed within each RBD family for the 14 families containing at least 10 proteins, sorted by effect-size magnitude. Markers denote the family-level point estimate; horizontal whiskers denote percentile-based 95% confidence intervals from 2,000 bootstrap resamples of proteins within each family; arrows indicate confidence intervals extending beyond the plotted range; n indicates the number of proteins in each family. Green markers correspond to positive point estimates and the red marker to the single family with a negative point estimate (PUA); the dashed vertical line marks d = 0.

## Discussion

In this study, we have presented CORAL, a deep learning model that integrates protein and RNA language models through a bidirectional cross-attention mechanism, and demonstrated two main contributions to the field of RNA–protein interaction. First, we establish that CORAL exhibits markedly superior generalization to novel RNA–protein pairs compared to existing methods, maintaining reasonable predictive performance under stringent non-redundancy conditions where competing approaches fall short. Second, we provide evidence that the cross-attention mechanism learns biologically meaningful representations, with specific attention heads systematically attending to structurally defined contact positions between RNA and protein molecules. Together, these findings advance both the practical utility and mechanistic interpretability of computational RPI prediction. The dramatic performance differences observed across our three evaluation strategies expose a fundamental problem in how the field has assessed RPI prediction models. On conventional random splits, all evaluated methods achieved strong performance, with F1 scores ranging from 0.83 to 0.92. This apparent success, however, masks a troubling reality: these benchmarks permit substantial sequence similarity between training and test sets, enabling models to succeed through memorization of sequence patterns associated with known interactions rather than learning generalizable principles of molecular recognition. The moderate performance degradation observed on pairwise splits suggests that models retain some capacity for generalization when at least one component of a test pair is sufficiently dissimilar to training examples. However, the component-wise split reveals the true extent of the generalization challenge. When we enforce that neither the RNA nor protein component of any test pair shares significant sequence similarity with training examples, performance collapses for all models, with F1 scores dropping as low as 0.03 for some methods. These patterns suggest that existing models memorize sequence signatures of molecules present in training data rather than capturing the principles governing RNA–protein recognition. CORAL’s performance on the component-wise split, maintaining an F1 score of 0.65 compared to 0.47 for the next-best method retaining discriminative behavior (ZHMolGraph), reflects meaningful progress toward genuinely generalizable RPI prediction. We attribute this improvement to the architectural choice of explicitly modeling inter-molecular relationships through cross-attention, thus promoting learning of interaction-relevant features that transfer to novel molecular contexts. A complementary architectural choice that contributes to this improved generalization is the joint fine-tuning of both sequence encoders through LoRA, rather than their use as frozen feature extractors as in prior methods. Our ablation analysis showed that adapting the encoder weights to the interaction prediction objective yields only modest gains on the redundant and pairwise splits, but produces a substantial improvement on the component-wise split. This pattern suggests that frozen pretrained representations, while adequate to support predictions when training and test sequences are similar, do not natively encode the features required for generalization. Nevertheless, an F1 score of 0.65 represents substantial room for improvement, and we emphasize that this result should be interpreted as evidence that the generalization problem in RPI prediction is genuinely difficult. Beyond predictive performance, our attention analysis provides insight into what CORAL has learned about RNA–protein recognition. The finding that specific attention heads, particularly heads 0 and 7, systematically exhibit elevated attention at structurally defined contact positions suggests that the model has discovered a correspondence between sequence-level features and three-dimensional binding interfaces. This effect is consistent across 79% of analyzed complexes. Conversely, 21% of complexes did not exhibit elevated attention at contact positions, indicating that while attention to structural contacts represents a primary strategy, the model likely employs multiple complementary mechanisms: sequence motifs, binding mode variations, or other features that override proximity-based signals in specific molecular contexts. A complementary line of attention evidence is provided by the protein-side, domain-resolution analysis (Figure 6). Marginalized over the RNA axis, CORAL’s cross-attention is elevated at residues falling within annotated RNA-binding domains across 94% of the 462 proteins examined (median within-protein Cohen’s *d* ≈ 0.94). The two analyses measure different facets of the same phenomenon at different resolutions: the contact analysis tests for elevated attention at specific atomic contact positions in 309 experimentally resolved complexes, while the domain analysis tests for elevated attention across whole 70–100-residue RBD modules in a much larger protein cohort. Together, the two analyses converge on the conclusion that CORAL’s cross-attention has internalized biologically meaningful features of RNA–protein recognition at multiple structural scales. Our study contributes not only a predictive model but also an evaluation framework that we believe merits adoption as a field standard. The distinction between pairwise and component-wise redundancy addresses a conceptual gap in how data leakage is typically conceived for interaction prediction tasks. Pairwise redundancy control, while preventing identical or near-identical interaction pairs from appearing in both training and test sets, permits a scenario in which a test RNA has been seen with different proteins during training, and vice versa. Under such conditions, a model might achieve high test performance by recognizing individual molecules rather than learning principles of their interaction. Component-wise redundancy control eliminates this possibility, requiring that predictions involve genuinely novel molecular entities. In conclusion, this work demonstrates that rigorous evaluation fundamentally changes our understanding of RPI prediction model capabilities. The apparent success of existing methods on conventional benchmarks reflects memorization rather than generalization. CORAL’s superior performance under stringent non-redundancy conditions, combined with interpretable attention patterns that correlate with structural binding interfaces, represents progress toward models that are both more reliable and more mechanistically transparent. As language model representations continue to improve for both proteins and nucleic acids, architectures that explicitly model cross-modal correspondence offer a promising framework for capturing the principles that govern biomolecular recognition.

## Materials and Methods

### Dataset sources

The positive RNA–protein interaction (RPI) samples used to construct the training and test sets for benchmarking CORAL against previously published methods were obtained from five publicly available databases: RNAInter, RPI488, RPI369, RPI2241, and RPI1807^29,41–43^. The RPI datasets comprise both interacting (positive) and non-interacting (negative) RNA–protein pairs derived from experimentally determined PDB complexes, with interactions defined based on distance criteria. For the present study, only positive pairs were retained from these datasets. RNAInter represents the largest RNA–protein interaction repository currently available, containing interactions curated from diverse sources including high-throughput experiments, low-throughput experiments, and computational predictions. However, only a subset of these interactions is supported by strong experimental evidence. The final positive dataset was therefore constructed by combining this high-confidence subset from RNAInter with the positive interactions from the four RPI datasets. Additionally, we excluded proteins exceeding 1,250 amino acid residues and RNAs exceeding 6,000 nucleotides in length, constraints imposed by GPU memory limitations during model training.

### Sequence clustering

To control redundancy levels within the dataset, RNA and protein sequences were clustered based on sequence identity using MMseqs2, an open-source software package implemented in C++ and optimized for searching and clustering large protein and nucleotide sequence sets^44^. The MMseqs2 “cluster” function was employed to generate protein sequence clusters at 30% and 60% sequence identity thresholds, and RNA sequence clusters at 50% and 80% sequence identity thresholds. The coverage parameter was fixed at 60% for all clustering operations, while remaining parameters were maintained at their default values.

### Pairwise split generation

To generate pairwise splits, interaction clusters were first defined by concatenating the cluster identifiers of the RNA and protein components for each RPI pair in the positive dataset, using the clusters obtained from MMseqs2. Since multiple RPI pairs may contain sequences belonging to identical RNA and protein clusters, duplicate interaction cluster identifiers were removed to obtain a set of unique interaction clusters. Subsequently, 20% of the unique interaction clusters were randomly assigned to the test set and 80% to the training set. The original RPI pairs were then partitioned according to their corresponding interaction cluster assignments. This procedure ensures that RPI pairs in the training and test sets belong to distinct interaction clusters, thereby preventing any test pair from sharing both RNA and protein cluster membership with any training pair (Figure 2b). Due to the stochastic nature of the cluster assignment process, training and test sets of varying sizes may result from different random seeds. To obtain splits with balanced dimensions, 100,000 independent random partitions were generated using different seeds. From these, the five splits satisfying the criterion of at least 12,000 positive interactions in the training set while maximizing the number of interactions in the test set were selected. This procedure was performed separately using protein and RNA clusters at 30% and 50% sequence identity, respectively, and at 60% and 80% sequence identity, respectively, yielding five splits for each threshold pair with approximately equivalent numbers of RPI pairs. Following the positive pair assignment, negative pairs were generated independently for the training and test sets through random pairing of non-interacting RNA and protein sequences (i.e., combinations absent from the positive dataset). The number of negative pairs was set equal to the number of positive pairs in each set, resulting in balanced datasets.

### Component-wise split generation

The component-wise split generation followed a similar workflow to the pairwise approach, with the key distinction that partitioning was based directly on individual RNA and protein clusters rather than on combined interaction clusters. Specifically, 20% of the protein clusters and 20% of the RNA clusters were independently and randomly assigned to the test set, with the remaining 80% assigned to the training set. The original RPI pairs were then partitioned based on the cluster assignments of their constituent sequences. Only RPI pairs for which both the RNA and protein components belonged to training-assigned clusters were included in the training set; conversely, only pairs for which both components belonged to test-assigned clusters were included in the test set. Pairs with mixed cluster assignments (i.e., one component in a training cluster and the other in a test cluster) were excluded from both sets (Figure 2b). This approach guarantees complete separation between training and test sets at the individual component level, such that no RNA or protein sequence in the test set shares cluster membership with any sequence in the training set. As with the pairwise splits, 100,000 independent random partitions were generated, and the five optimal splits were selected based on the same criteria. Negative pair generation was performed identically to the pairwise split procedure, maintaining a 1:1 ratio of positive to negative pairs.

### CORAL architecture

CORAL utilizes LLMs DNABERT2 and ESM2 models as the sequence encoders for RNA and Protein sequences respectively. These BERT-like models were pretrained to generate information rich 768-dimensional embeddings for RNAs and 640-dimensional embeddings for proteins. Both the LLMs models are fine tuned with Low-Rank Adaptation (LoRA) during CORAL’s training.

### LLMs embedding layer

CORAL employs pre-trained large language models (LLMs) as modality-specific sequence encoders to extract contextualized token-level representations for RNA and protein inputs. RNA sequences are processed by a transformer-based nucleotide language model that generates embeddings with a hidden dimensionality of 768. Protein sequences are encoded using ESM2, a state-of-the-art protein language model pre-trained on evolutionary-scale sequence data, which produces token embeddings with a native hidden dimensionality of 640. Given a nucleotide sequence *x*_0_ and an amino acid sequence *x*_1_, we encode them using DNABERT2 and ESM2:

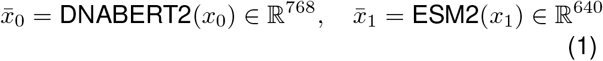

where 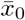 and 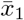 are the LLMs embeddings for *x*_0_ and *x*_1_. Both encoders are initialized from pre-trained checkpoints via the HuggingFace Transformers library and are fine tuned with Low-Rank Adaptation (LoRA) during CORAL’s training.

### Cross-modality module

The cross-modality module serves as the central architectural component of CORAL, implementing a bidirectional cross-attention mechanism to establish explicit inter-modal correspondence between RNA and protein representation spaces. Prior to cross-modal fusion, the protein token embeddings obtained from ESM2 are transformed through a learned linear projection layer that maps the 640-dimensional protein representations to a 768-dimensional space, thereby ensuring dimensional congruence with the RNA embeddings, a prerequisite for the computation of cross-attention scores without inducing dimension mismatch errors. The cross-modality encoder comprises *N* stacked cross-modality layers (default *N* = 2), each executing three sequential computational stages. The initial stage implements bidirectional cross-attention through two parallel attention blocks with independent parameterization: an RNA-to-protein attention block wherein RNA token embeddings constitute the query vectors while protein token embeddings serve as keys and values, and a symmetric protein-to-RNA attention block operating in the opposite direction. This bidirectional formulation enables each modality to selectively aggregate contextually relevant features from the complementary sequence representation. Subsequently, modality-specific self-attention blocks are applied independently to the RNA and protein feature streams, permitting intra-sequence contextual refinement while preserving modality-specific representational structure. The terminal stage of each layer comprises position-wise feed-forward networks (FFNs) with GELU non-linearity and an intermediate dimensionality expansion factor of four, applied independently to each modality stream. In accordance with standard transformer methodology, residual connections followed by layer normalization are incorporated after each attention sub-layer and FFN to promote gradient propagation and stabilize optimization dynamics. All attention computations utilize multi-head attention with 12 parallel attention heads and a per-head dimensionality of 64.

### Pooling and output heads

Aggregation of variable-length token-level representations into fixed-dimensional sequence embeddings is achieved through a pooling operation applied to the outputs of the cross-modality encoder. Two pooling strategies were evaluated: extraction of the [CLS] token representation and mean pooling over non-padding token positions. Mean pooling, implemented as the attention-mask-weighted average of token embeddings with numerical stability provisions for variable sequence lengths, was selected based on superior empirical performance. The resultant RNA and protein sequence embeddings are concatenated to yield a joint representation of dimensionality 1536, which is subsequently projected to a 768-dimensional latent space via a linear transformation followed by hyperbolic tangent activation. The architecture incorporates three task-specific output heads within a multi-task learning framework. Two masked language modeling (MLM) heads, dedicated to RNA and protein modalities respectively, operate on the token-level representations to reconstruct masked input tokens during pre-training. Each MLM head comprises a linear transformation, GELU activation, layer normalization, and a terminal projection layer whose weights are tied to the corresponding encoder’s input embedding matrix, following standard practice for language model pre-training objectives. The interaction prediction head consists of a linear classifier that receives the pooled joint embedding and produces logits for binary classification of RNA–protein interaction probability.

### Model training and evaluation

CORAL was trained from scratch using a multi-task learning framework that jointly optimized two complementary objectives. The first objective was masked language modeling, in which 15% of tokens in both RNA and protein sequences were randomly masked and reconstructed through the cross-attention encoder, encouraging the development of robust contextual representations of sequence patterns. The second objective was binary cross-entropy classification for discriminating between interacting and non-interacting RNA–protein pairs. The composite training objective was computed as an unweighted sum of the two loss terms. Low-Rank Adaptation (LoRA) with a rank of 8 was employed for parameter-efficient training of the two sequence encoders, while the cross-modality module and output heads were fully trainable. Optimization was performed using AdamW with default hyperparameters coupled with a linear learning rate scheduler with a warm-up phase corresponding to 5% of total training step. The model was trained for 10 epochs with a batch size of 8 on the benchmark data partitions described in Dataset Sources. For the LoRA-fine-tuned variant, the initial learning rate was set to 1 *×* 10^−6^, a value chosen to permit gradual adaptation of the pretrained encoder weights while preserving the representations acquired during large-scale pretraining. For the frozen-encoder ablation, in which only the cross-modality encoder, the protein projection layer, and the classification head were trainable, the initial learning rate was raised to 1 *×* 10^−4^ to enable effective optimization of these randomly initialized components, which would otherwise have remained substantially under-trained within the same epoch budget. This choice was conservative with respect to our conclusions, as it provided the frozen-encoder baseline with a more favorable optimization regime.

### Statistical analysis of attention patterns

To assess whether the cross-attention mechanism systematically prioritizes structurally defined contact positions, we employed a hierarchical analytical framework comprising three complementary statistical approaches. Throughout this analysis, the per-head attention signal was defined as the geometric mean of the two directional maps produced by the final cross-modality layer, 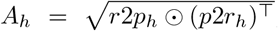, where *r*2*p*_*h*_ is head h’s RNA-to-protein attention map, (*p*2*r*_*h*_)^⊤^ is the transpose of its protein-to-RNA map aligned to the same RNA-by-protein grid, and ⊙ denotes the element-wise (Hadamard) product. This symmetric combination reflects the direction-independent nature of a physical contact, which is registered regardless of read-out direction. The two heads on which the contact analyses were centred (heads 0 and 7) were selected by a prior screen across all 12 heads, a Friedman omnibus test followed by post-hoc Benjamini–Hochberg-corrected Wilcoxon signed-rank comparisons against the head with the largest median per-complex Δ, mirroring the median-based reference-head criterion used in the domain-resolution analysis. All contact statistics described below were computed on *A*_*h*_ (for the screened-head analyses, on the average of *A*_0_ and *A*_7_). A critical consideration underlying our methodology was the nested structure of the data: attention scores within each RNA–protein complex are not independent observations, as they derive from the same attention matrix and share complex-specific properties such as sequence length, binding mode, and contact density. Treating individual position pairs as independent observations would constitute pseudoreplication, artificially inflating sample sizes and yielding spuriously significant results. To address this concern, we treated each complex as the independent unit of analysis throughout all statistical procedures, computing summary statistics at the complex level before performing population-level inference. For the primary meta-analytic approach, we computed the mean difference between contact and non-contact attention scores for each complex (Δ_*i*_ = mean contact attention − mean non-contact attention), yielding 309 independent observations. To test whether the population-level mean difference significantly deviated from zero, we performed a one-sample *t*-test on these per-complex differences. To complement the *t*-test with an approach that makes minimal distributional assumptions, we applied the non-parametric sign test. This test evaluates whether the proportion of complexes exhibiting positive differences significantly exceeds the chance expectation of 50%, relying solely on the direction rather than the magnitude of each observation. The sign test assumes only that observations are independent and drawn from a continuous distribution; it makes no assumptions regarding distributional shape, variance homogeneity, or the absence of outliers. This property renders it particularly valuable for validating findings when the distribution of effect sizes may be skewed or heavy-tailed, as can occur in biological datasets with heterogeneous molecular contexts. Furthermore, by assessing directional consistency rather than mean magnitude, the sign test provides an independent measure of whether the observed bias generalizes across the majority of complexes rather than being driven by a subset of extreme values. To further validate our findings under relaxed distributional assumptions and obtain robust confidence intervals, we performed hierarchical bootstrap resampling. Critically, our implementation respected the hierarchical structure of the data by resampling at the level of complexes rather than individual position pairs, thereby preserving the dependence structure within each attention matrix. In each of 10,000 iterations, we resampled 309 complexes with replacement and computed the mean Δ_*i*_ for the re-sampled population. This procedure yields an empirical sampling distribution from which we derived percentile-based 95% and 99% confidence intervals. The bootstrap assumes that the observed sample is representative of the underlying population but makes no parametric assumptions regarding the shape of the distribution, providing a robust complement to the *t*-test that accounts for potential non-normality and inter-complex heterogeneity. To move beyond statistical significance and quantify effect magnitude at the individual complex level, we computed Cohen’s *d* independently for each of the 309 complexes. Cohen’s *d* is a standardized effect size metric defined as the difference between two group means divided by the pooled standard deviation, expressing the magnitude of the difference in units of standard deviation. This standardization enables meaningful comparison of effect sizes across complexes that differ in their underlying variance structures, which is essential given the heterogeneity in sequence lengths and contact densities across our dataset. By examining the distribution of these 309 effect sizes, we could assess whether any observed population-level bias reflects a consistent pattern across individual complexes or represents an aggregation artifact driven by a small number of outliers with disproportionately large effects. Finally, to investigate functional differentiation among attention heads and validate our focus on heads 0 and 7, we repeated all three analytical approaches using attention scores averaged across all twelve heads in the cross-attention mechanism. This comparative analysis enabled assessment of whether any observed contact-position bias represents a specialized property of specific heads or a general feature distributed across the attention mechanism. A substantial attenuation of the effect when averaging across all heads would suggest functional specialization, wherein different heads encode distinct and potentially complementary aspects of RNA–protein interactions. All statistical analyses were conducted in Python (v3.10) using SciPy (v1.15.3) for parametric and non-parametric tests. Statistical significance was assessed at *α* = 0.05 unless otherwise specified.

### Analysis of attention at RNA-binding domains

To assess whether CORAL’s protein-side cross-attention is systematically elevated at residues annotated as RNA-binding domains, we conducted a domain-level analog of the contact-position analysis on a substantially larger protein cohort drawn from the curated interaction dataset. The RBD whitelist was constructed from the human RBP census of Gerstberger and colleagues^6^. Among the RBD families catalogued in that census, we retained those observed in at least four distinct RBPs, yielding 85 families spanning canonical sequence-specific RBDs (including RRM, KH, dsRBD, CCCH and CCHC zinc fingers, CSD, PUF, Piwi, YTH, La, Sm/LSM, S1, R3H, SAP, PWI, Surp, and TUDOR) together with RNA-associated modules (ribosomal proteins, translation factors, DEAD-box helicases, GTPases, tRNA synthetases, and RNA-modification and -processing enzymes). The Pfam accession identifier was retrieved for each of the 85 families. The full set of proteins appearing as the positive partner in any interaction in the curated dataset was then annotated against Pfam using hmmscan from the HMMER suite, retaining only hits matching one of the 85 whitelisted accessions. Within each protein, overlapping or adjacent Pfam hits were merged into non-overlapping intervals, and the per-protein RBD residue mask was defined as the union of these intervals across all annotated RBDs. Protein-level quality filters were applied to ensure that the inside-versus-outside contrast was statistically meaningful. We retained only proteins that (i) carried at least one annotated RBD interval, (ii) exceeded 100 amino acid residues in total length, (iii) exhibited RBD coverage not exceeding 70% of the total length, and (iv) had at least one true-positive RNA partner, defined as a curated-positive interaction predicted by CORAL with probability greater than 0.5 at a 0.5 decision threshold. The application of these criteria yielded 462 proteins aggregated from 12,834 true-positive (protein, RNA) pairs. For each (protein, RNA) pair in the analysis set, inference was performed with CORAL and the RNA-to-protein cross-attention map was extracted from the final cross-modality layer and averaged over the selected attention head(s). The resulting two-dimensional map (RNA tokens by protein residues) was then marginalized over the RNA axis by mean reduction to obtain a per-residue protein attention score; mean rather than sum reduction was chosen to length-normalize the per-residue score and avoid biasing it toward proteins observed with longer RNA partners. The per-pair vectors were subsequently averaged element-wise across each protein’s true-positive partners to produce a per-protein consensus attention profile. Under this aggregation scheme each protein constitutes one independent observation in subsequent statistical tests, paralleling the per-complex unit of analysis used for the contact-position analysis and avoiding the pseudoreplication that would arise from treating individual (protein, RNA) pairs as independent units. To identify which of the 12 cross-attention heads exhibit differential responses at RBD residues while controlling for multiple comparisons, we adapted the head-screening procedure used for the contact analysis. For each protein and each head *h*, we computed the per-head difference Δ_*h*_ = mean(attention | residue ∈ any RBD) − mean(attention | residue ∉ any RBD) on the protein-marginalized scores. A Friedman test was applied across the 12 heads on the resulting 462 × 12 matrix of per-protein per-head differences. Conditional on a significant Friedman result, we identified the head with the largest median Δ_*h*_ across proteins and performed post-hoc pairwise Wilcoxon signed-rank tests comparing it against each of the remaining heads, with multiple-comparison correction by the Benjamini– Hochberg false discovery rate procedure at *α* = 0.05. Heads not significantly different from the best-ranked head would have been retained alongside it; in our analysis no other head met this criterion, and head 2 alone was selected for the main test. The same screening procedure was applied independently from the head selection of the contact-position analysis to allow for the possibility that distinct attention heads specialize for distinct interpretability signals. For each protein included in the analysis we then computed Δ_*k*_ = mean(*s* | residue ∈ any RBD) − mean(*s* | residue ∉ any RBD), where s is the per-protein consensus attention vector. Proteins were required to contribute at least three residues to each of the inside and outside sets; this condition was satisfied by all 462 filtered proteins. We applied four complementary statistical approaches treating the protein as the independent unit of analysis. First, a one-sample t-test (alternative = greater) was performed on the population of Δ_*k*_. Second, an exact one-sided binomial sign test assessed whether the proportion of proteins with Δ_*k*_ *>* 0 exceeded the chance expectation of 50%. Third, a hierarchical bootstrap with 10,000 iterations (random seed 42) was conducted by resampling proteins, rather than individual residues, with replacement, yielding the bootstrap standard error and the percentile-based 95% confidence interval for the population mean Δ_*k*_. Fourth, the per-protein Cohen’s d was computed as 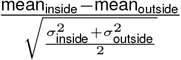, with *σ*^2^ denoting the population variance within each residue set; this within-protein effect size quantifies the standardized inside-versus-outside separation for an individual protein, and is summarized across the cohort as a distribution. To test whether the observed bias depends on the specific placement of RBD intervals along the protein sequence rather than reflecting a generic positional confound, we constructed a position-shuffle null distribution. For each protein, the binary RBD residue mask was cyclically rotated along the sequence by a random non-zero integer offset; this operation preserves the number, lengths, and gap structure of the annotated intervals while randomizing their absolute placement. The aggregate mean Δ was recomputed under the rotated masks and the procedure repeated 1,000 times (random seed 42). The empirical p-value was defined as the proportion of null draws yielding an aggregate Δ greater than or equal to the observed value. To test whether the population-level effect generalized across diverse RBD classes, proteins were grouped by their dominant annotated RBD family, with dominance assigned by the largest cumulative annotated interval length when more than one family was present. Families containing at least 10 proteins were retained, yielding 14 strata. Within each family, a one-sample t-test (alternative = greater) was performed on the per-protein Δ_*k*_, and an across-protein effect size was computed as mean(Δ_*k*_) / SD(Δ_*k*_) using the sample standard deviation. Percentile-based 95% confidence intervals for this across-protein effect size were obtained by resampling proteins within the family with replacement over 2,000 bootstrap iterations (random seed 43). This across-protein effect size, visualized as a forest plot in Figure 6d, is a distinct construct from the within-protein pooled Cohen’s d used in the main test and visualized as the cumulative distribution function in Figure 6c, and the two are not directly comparable in magnitude. All statistical computations were performed in Python (v3.10) using SciPy (v1.15.3) for parametric and non-parametric tests, consistent with the implementation used for the contact-position analysis. Statistical significance was assessed at *α* = 0.05 unless otherwise specified.

## Resource availability

### Lead contact

Requests for further information and resources should be directed to and will be fulfilled by the lead contact, Pier Federico Gherardini.

### Materials availability

This study did not generate new unique reagents.

### Data and code availability

- Processed data and the code used for analysis are available on GitHub at https://github.com/MarioCatalano99/Coral. Additional details and usage instructions are provided in the repository README file.
- Any additional information required to reanalyze the data reported in this paper is available from the lead contact upon request.

## Author contributions

**Conceptualization:** P.F.G., M.H.-C.

**Methodology:** M.C.

**Software:** M.C.

**Validation:** M.C.

**Formal Analysis:** M.C.

**Investigation:** M.C.

**Resources:** M.C.

**Data Curation:** M.C., R.A.

**Writing – Original Draft:** M.C.

**Writing – Review & Editing:** M.C., P.F.G., C.M.W., G.G., G.P., M.H.-C., G.A.

**Visualization:** M.C.

**Supervision:** P.F.G., G.G., M.H.-C.

**Funding Acquisition:** M.H.-C.

## Acknowledgements

This research was funded by the European Union—NextGenerationEU: National Center for Gene Therapy and Drugs based on RNA Technology, CN3—Spoke 7 (code: CN00000041; PNRR—Mission 4, Component 2; Investment 1.4)

## Supplementary Information

**Supplementary Figure 1.**
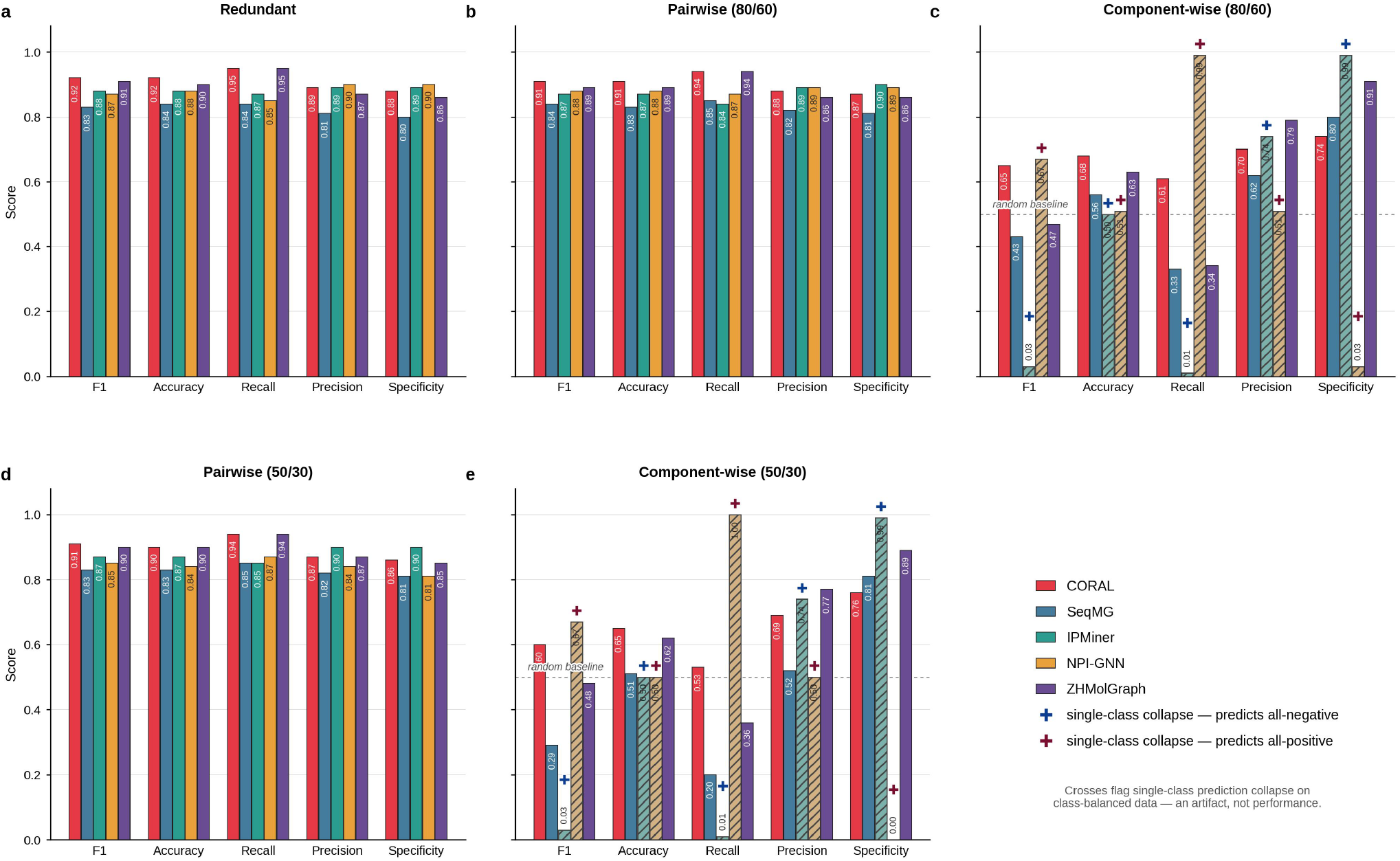
Full metric breakdown for CORAL and baseline methods across all five data partitions. F1, Accuracy, Recall, Precision and Specificity are shown for five RNA–protein interaction predictors: CORAL (this work, red), SeqMG (blue), IPMiner (teal), NPI-GNN (amber) and ZHMolGraph (purple), on **(a)** the conventional redundant split, **(b)** the pairwise (80/60) non-redundant split, **(c)** the component-wise (80/60) non-redundant split, **(d)** the pairwise (50/30) non-redundant split, and **(e)** the component-wise (50/30) non-redundant split. Numbers in parentheses denote the sequence identity thresholds (RNA / protein) used to construct each partition; the redundant split is threshold-independent. Bar heights are means over five independent train/test folds and the exact value is annotated on each bar. The dashed grey reference line in (c) and (e) marks the score expected from a random predictor on the class-balanced (50:50 positives:negatives) test sets. Crosses flag methods that collapse to single-class prediction on the component-wise splits; for these methods, F1, Recall and Specificity are mechanically determined by the direction of collapse and the class balance and therefore do not reflect discriminative ability. Cross colour encodes the direction of collapse: navy denotes “predicts all-negative”, whereas maroon denotes “predicts all-positive”. Diagonal hatching and desaturated colour of the affected bars visually reinforce these crosses. Panels (d) and (e) show that the performance degradation reported in the main text intensifies under the stricter identity thresholds (50% RNA, 30% protein), with CORAL retaining the highest F1 and Accuracy on the component-wise split while the competing methods degrade further.

